# A validation of the diathesis-stress model for depression in Generation Scotland

**DOI:** 10.1101/389494

**Authors:** Aleix Arnau-Soler, Mark J. Adams, Toni-Kim Clarke, Donald J. MacIntyre, Keith Milburn, Lauren Navrady, Generation Scotland, Major Depressive Disorder Working Group of the Psychiatric Genomics Consortium, Caroline Hayward, Andrew McIntosh, Pippa A. Thomson

**Affiliations:** Medical Genetics Section, University of Edinburgh, Centre for Genomic and Experimental Medicine and MRC Institute of Genetics and Molecular Medicine, Edinburgh, UK; Division of Psychiatry, Deanery of Clinical Sciences, University of Edinburgh, Royal Edinburgh Hospital, Morningside Park, Edinburgh EH10 5HF, UK; Health Informatics Centre, University of Dundee, UK; A collaboration between the University Medical School and NHS in Aberdeen, Dundee, Edinburgh and Glasgow, Scotland, UK; For a full list of MDD working group of the PGC investigators, see the Supplementary Material; Medical Research Council Human Genetics Unit, Institute of Genetics and Molecular Medicine, University of Edinburgh, Edinburgh, UK; Centre for Cognitive Ageing and Cognitive Epidemiology, University of Edinburgh, Edinburgh, UK

## Abstract

Depression has well-established influences from genetic and environmental risk factors. This has led to the *diathesis-stress* theory, which assumes a multiplicative gene-by-environment interaction (GxE) effect on risk. Recently, *Colodro-Conde et al*. empirically tested this theory, using the polygenic risk score for major depressive disorder (PRS, genes) and stressful life events (SLE, environment) effects on depressive symptoms, identifying significant GxE effects with an additive contribution to liability. We have tested the *diathesis-stress* theory on an independent sample of 4 919 individuals.

We identified nominally significant positive GxE effects in the full cohort (R^2^ = 0.08%, *p* = 0.049) and in women (R^2^ = 0.19%, *p* = 0.017), but not in men (R^2^ = 0.15%, *p* = 0.07). GxE effects were nominally significant, but only in women, when SLE were split into those in which the respondent plays an active or passive role (R^2^ = 0.15%, *p* = 0.038; R^2^ = 0.16%, *p* = 0.033, respectively). High PRS increased the risk of depression in participants reporting high numbers of SLE (*p* = 2.86 × 10^−4^). However, in those participants who reported no recent SLE, a higher PRS appeared to increase the risk of depressive symptoms in men (β = 0.082, *p* = 0.016) but had a protective effect in women (β = −0.061, *p* = 0.037). This difference was nominally significant (*p* = 0.017). Our study reinforces the evidence of additional risk in the aetiology of depression due to GxE effects. However, larger sample sizes are required to robustly validate these findings.

## INTRODUCTION

Stressful life events (SLE) have been consistently recognized as a determinant of depressive symptoms, with many studies reporting significant associations between SLE and major depressive disorder (MDD)^1-7^. Some studies suggest that severe adversity is present before the onset of illness in over 50% of individuals with depression^8^ and may characterize a subtype of cases^9^. However, some individuals facing severe stress never present symptoms of depression^10^. This has led to a suggestion that the interaction between stress and an individual’s vulnerability, or *diathesis*, is a key element in the development of depressive symptoms. Such vulnerability can be conceived as a set of biological factors that predispose to illness. Several *diathesis-stress* models have been successfully applied across many psychopathologies^11-15^.

The *diathesis-stress* model proposes that a latent *diathesis* may be activated by stress before psychopathological symptoms manifest. Some levels of *diathesis* to illness are present in everybody, with a threshold over which symptoms will appear. Exceeding such a threshold depends on the interaction between *diathesis* and the degree of adversity faced in SLE, which increases the liability to depression beyond the combined additive effects of the *diathesis* and stress alone^11^. Genetic risk factors can, therefore, be conceived as a genetic *diathesis*. Thus, this genetically driven effect produced by the *diathesis-stress* interaction can be seen as a gene-by-environment interaction (GxE).

MDD is characterized by a highly polygenic architecture, composed of common variants with small effect and/or rare variants^16^. Therefore, interactions in depression are also expected to be highly polygenic. In recent years, with the increasing success of genome-wide association studies, GxE studies in depression have shifted towards hypothesis-free genome-wide and polygenic approaches that capture liability to depression using genetic data^17-25^. Recent advances in genomics and the massive effort from national institutions to collect genetic, clinical and environmental data on large population-based samples now provide an opportunity to empirically test the *diathesis-stress* model for depression. The construction of polygenic risk scores (PRS) offers a novel paradigm to quantify genetic *diathesis* into a single genetic measure, allowing us to study GxE effects with more predictive power than any single variant^26-29^. PRS are genetic indicators of the aggregate number of risk alleles carried by an individual weighted by their allelic effect estimated from genome-wide association studies. This polygenic approach to assessing the *diathesis-stress* model for depression has been tested using either childhood trauma^17,19,25^ or adult SLE^18,23,25^ as measures of environmental adversity.

Recently, Colodro-Conde *et al*.^23^ provided a direct test of the *diathesis-stress* model for recent SLE and depressive symptoms. In this study, Colodro-Conde *et al*. used PRS weighted by the most recent genome-wide meta-analysis conducted by the Psychiatric Genetics Consortium (PGC; N = 159 601), and measures of three environmental exposures: lack of social support, “personal” SLE, and “network” SLE. Colodro-Conde *et al*. reported a significant additive risk on liability to depression due to a GxE effect in individuals who combine a high genetic predisposition to MDD and a high number of reported “personal” SLE, mainly driven by effects in women. A significant effect of interaction was not detected in males. They found no significant interaction between the genetic *diathesis* and “network” SLE or social support. They concluded that the effect of stress on risk of depression was dependent on an individual’s *diathesis*, thus supporting the *diathesis-stress* theory. In addition, they suggested possible sex-specific differences in the aetiology of depression. However, Colodro-Conde *et al*. findings have not, to our knowledge, been independently validated.

In the present study we aim to test the *diathesis-stress* model in an independent sample of 4 919 unrelated white British participants from a further longitudinal follow-up from Generation Scotland and assess the differences between women and men, using self-reported depressive symptoms and recent SLE.

## MATERIALS AND METHODS

### Sample description

Generation Scotland is a family-based population cohort recruited throughout Scotland by a cross-disciplinary collaboration of Scottish medical schools and the National Health Service (NHS) between 2006 and 2011^30^. At baseline, blood and salivary DNA samples from Generation Scotland participants were collected, stored and genotyped at the Wellcome Trust Clinical Research Facility, Edinburgh. Genome-wide genotype data were generated using the Illumina HumanOmniExpressExome-8 v1.0 DNA Analysis BeadChip (San Diego, CA, USA) and Infinium chemistry^31^. The procedures and further details for DNA extraction and genotyping have been extensively described elsewhere^32,33^. In 2014, 21 525 participants from Generation Scotland eligible for re-contact were sent self-reported questionnaires as part of a further longitudinal assessment funded by a Wellcome Trust Strategic Award “STratifying Resilience and Depression Longitudinally” (STRADL)^34^ to collect new and updated mental health questionnaires including psychiatric symptoms and SLE measures. 9 618 re-contacted participants from Generation Scotland agreed to provide new measures to the mental health follow-up^34^ (44.7% response rate). Duplicate samples, those showing sex discrepancies with phenotypic data, or that had more than 2% missing genotype data, were removed from the sample, as were samples identified as population outliers in principal component analysis (mainly non-Caucasians and Italian ancestry subgroup). In addition, individuals with diagnoses of bipolar disorder, or with missing SLE data, were excluded from the analyses. SNPs with more than 2% of genotypes missing, Hardy-Weinberg Equilibrium test *p* < 1 × 10^−6^, or a minor allele frequency lower than 1%, were excluded. Individuals were then filtered by degree of relatedness (pi-hat < 0.05) using PUNK v1.9^35^, maximizing retention of those participants reporting higher numbers of SLE (see phenotype assessment below). After quality control, the final dataset comprised 4 919 unrelated individuals of European ancestry and 560 351 SNPs (mean age at questionnaire: 57.2, s.d. = 12.2, range 22-95; *women*: *n* = 2 990 - 60.8%, mean age 56.1, s.d. = 12.4; *men*: *n* = 1 929 - 39.2%, mean age 58.7, s.d. = 11.8). Further details on the recruitment procedure and Generation Scotland profile are described in detail elsewhere^30,32,36-38^. All participants provided written consent. All components of Generation Scotland and STRADL obtained ethical approval from the Tayside Committee on Medical Research Ethics on behalf of the National Health Service (reference 05/s1401/89). Generation Scotland data is available to researchers on application to the Generation Scotland Access Committee.

### Phenotype assessment

Participant self-reported current depressive symptoms through the 28-item scaled version of The General Health Questionnaire^39,40^. The General Health Questionnaire is a reliable and validated psychometric screening tool to detect common psychiatric and non-psychotic conditions (General Health Questionnaire Cronbach alpha coefficient: 0.82 – 0.86)^41^. This consists of 28 items designed to identify whether an individual’s current mental state has changed over the last 2 weeks from their typical state. The questionnaire captures core symptoms of depression through subscales for severe depression, emotional (e.g. anxiety and social dysfunction) and somatic symptoms linked to depression. These subscales are highly correlated^42^ and suggest an overall general factor of depression^43^. Participants rated the 28 items on a four-point Likert scale from 0 to 3 to assess its degree or severity^41^ (e.g., *Have you recently felt that life is entirely hopeless?* “Not at all”, “No more than usual”, “Rather more than usual”, “Much more than usual”), resulting on an 84-point scale depression score. The Likert scale, which provides a wider and smoother distribution^41^, could be more sensitive to detect changes in mental status in those participants with chronic conditions or chronic stress who may feel their current symptoms as “usual^-44^, and to detect psychopathology changes as response to stress. The final depression score was log transformed to reduce the effect of positive skew and provide a better approximation to a normal distribution. In addition, participants completed the Composite International Diagnostic Interview–Short Form, which diagnoses lifetime history of MDD according to DSM-IV criteria^45^. The depression score predicted lifetime history of MDD (odd ratio = 1.91, 95% confidence intervals 1.80-2.02, *p* = 1.55 × 10^−102^, N = 8 994), with a 3.8-fold increased odd of having a lifetime history of MDD between participants in the top and bottom deciles, thus supporting the usefulness of the depression score in understanding MDD. For a better interpretation, we scaled the depression score to a mean of 0 when required (Figure 3).

Data from a self-reported questionnaire based on the List of Threating Experiences^46^ was used to construct a measure of common SLE over the previous 6 months. The List of Threatening Experiences is a reliable psychometric device to measure psychological “stress^-47,48^. It consists of a 12-item questionnaire to assess SLE with considerable long-term contextual effects (e.g., *Over last 6 months, did you have a serious problem with a close friend, neighbour or relatives?)*. A final score reflecting the total number of SLE (TSLE) ranging from 0 to 12 was constructed by summing the “yes” responses. Additionally, TSLE was split into two categories based on those items measuring SLE in which the individual may play and active role exposure to SLE, and therefore in which the SLE is influenced by genetic factors and thus subject to be “dependent” on an individual’s own behaviour or symptoms (DSLE; 6 items, e.g., *a serious problem with a close friend, neighbour or relatives* may be subject to a respondent’s own behaviour), or SLE that are not influenced by genetic factors, likely to be “independent” on a participant’s own behaviour (ISLE; 5 items, e.g., *a serious illness, injury or assault happening to a close relative* is potentially independent of a respondent’s own behaviour)^46,49^. The item *“Did you/your wife or partner give birth?”* was excluded from this categorization. In addition, SLE reported were categorized to investigate the *diathesis* effect at different levels of exposure, including a group to test the *diathesis* effect when SLE is not reported. 3 levels of SLE reported were defined (0 SLE = “none”, 1 or 2 SLE = “low”, and 3 or more SLE = “high”) to retain a large enough sample size for each group to allow meaningful statistical comparison.

### Polygenic profiling & statistical analysis

Polygenic risk scores (PRS) were generated by PRSice^50^, whose functionality relies mostly on PLINK v1.9^35^, and were calculated using the genotype data of Generation Scotland participants (i.e. target sample) and summary statistics for MDD from the PGC-MDD2 GWAS release (July 2016, discovery sample) used by Colodro-Conde *et al*.^23^, with the added contribution from QIMR cohort and the exclusion of Generation Scotland participants, resulting in summary statistics for MDD derived from a sample of 50 455 cases and 105 411 controls.

Briefly, PRSice removed strand-ambiguous SNPs and clump-based pruned (r^2^ = 0.1, within a 10Mb window) our target sample to obtain the most significant independent SNPs in approximate linkage equilibrium. Independent risk alleles were then weighted by the allelic effect sizes estimated in the independent discovery sample and aggregated into PRS. PRS were generated for eight *p* thresholds (*p* thresholds: < 5 × 10^−8^, < 1 × 10^−5^, < 0.001, < 0.01, < 0.05, < 0.1, < 0.5, <=1) determined by the discovery sample and standardized (See Supplementary Table 1 for summary of PRS).

A genetic relationship matrix (GRM) was calculated for each dataset (i.e. *full cohort, women* and *men*) using GCTA1.26.0^51^. Mixed linear models using the GRM were used to estimate the variance in depression score explained by PRS, SLEs and their interaction; and stratified by sex. 20 principal components were calculated for the datasets.

The mixed linear model used to assess the effects of PRS is as follows:

*Depression* = *β*_0_ + *β*_1_*TPRS* + *GRM* + *Covariates*

Mixed linear models used to assess the effect of the stressors are as follows:

*Depression* = *β*_0_ + *β*_1_*TSLE* + *GRM* + *Covariates*
*Depression* = *β*_0_ + *β*_1_*DSLE* + *GRM* + *Covariates*
*Depression* = *β*_0_ + *β*_1_*ISLE* + *GRM* + *Covariates*

Following Colodro-Conde *et al*.^23^, (i.e. age, age^2^, sex, age-by-sex and age^2^- by-sex interactions, and 20 principal components) were regressed from PRS (PRS’) and SLE scores (i.e. TSLE’, DSLE’ and ISLE’; SLEs’) before fitting models in GCTA to guard against confounding influences on the PRS-by-SLEs interactions^52^. PRS’ and SLEs’ were standardized to a mean of 0 and a standard deviation of 1. The Mixed linear models (i.e. the *diathesis-stress* model) used to assess GxE effects are as follows:

*Depression* = *β*_0_ + *β*_1_*PRS*′ + *β*_2_*TSLE*′ + *β*_3_*PRS*′*xTSLE*′ + *GRM* + *Covariates*
*Depression* = *β*_0_ + *β*_1_*PRS*′ + *β*_2_*DSLE*′ + *β*_3_*PRS*′*xDSLE*′ + *GRM* + *Covariates*
*Depression* = *β*_0_ + *β*_1_*PRS*′ + *β*_2_ *ISLE*′ + *β*_3_*PRS*′*xISLE*′ + *GRM* + *Covariates*

Covariates fitted in the models above were age, age^2^, sex, age-by-sex, age^2^-by-sex and 20 principal components. Sex and its interactions (age-by-sex and age^2^-by-sex) were omitted from the covariates when stratifying by sex. All parameters from the models were estimated using GCTA and the significance of the effect (*β*) from fixed effects assessed using a Wald test. The significance of main effects (PRS and SLEs) allowed for nominally testing the significance of interactions at *p*-threshold = 0.05. To account for multiple testing correction, a Bonferroni’s adjustment correcting for 8 PRS and 3 measures of SLE tested (24 tests) was used to establish a robust threshold for significance at *p* = 2.08 × 10^−3^.

The PRS effect on depression score at different levels of exposure was further examined for the detected nominally significant interactions by categorizing participants on three groups based on the number of SLE reported (i.e. “none”, “low” or “high”). Using linear regression, we applied a least squares approach to assess PRS’ effects on the depression score in each SLE category. Further conservative Bonferroni correction to adjust for the 3 SLE categories tested established a threshold for significance of *p* = 6.94 × 10^−4^.

Differences on the estimated size of GxE effect between women and men were assessed by comparing a z-score to the standard normal distribution (*α* = 0.05, one-tailed). Z-scores were derived from GxE estimates (*β*) and standard errors (SE) detected in women and men as follows:

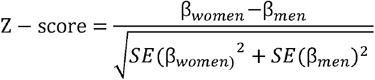

## RESULTS

PRS for MDD significantly predicted the depression score across the whole sample (β = 0.080, s.e. = 0.014, *p* = 7.53 × 10^−9^) explaining 0.64% of the variance at its best *p*-threshold (*p*-threshold = 0.1; Figure 1a). Stratifying by sex, PRS significantly predicted the depression score in both sexes, explaining 0.59% in men and 0.67% in women (*men*: *p*-threshold = 0.1, β = 0.077, s.e. = 0.022, *p* = 2.09 × 10^−4^; *women*: *p*-threshold = 0.1, β = 0.082, s.e. = 0.018, *p* = 4.93 × 10^−6^; Figure 1a). Self-reported SLE over the last 6 months (TSLE, mean = 1.3 SLE, s.d. = 1.5) also significantly predicted depression score for the whole sample and stratified by sex (*full cohort:* variance explained = 4.91%, β = 0.222, s.e. = 0.014, *p* = 9.98 × 10^−59^; *men:* 4.19%, β = 0.205, s.e. = 0.021, *p* = 2.23 × 10^−22^; *women:* 5.33%, β = 0.231, s.e. = 0.018, *p* = 7.48 × 10^−38^; Figure 1b). Overall, significant additive contributions from genetics and SLE in depression score were detected in all participants and across sexes. There was no significant difference in the direct effect of TSLE between women and men (*p* = 0.17). However, the variance in depression score explained by the TSLE appeared to be lower than the variance explained by the measure of personal SLE (PSLE) used in Colodro-Conde *et al*.^23^ (12.9%). This may, in part, be explained by different contributions of dependent and independent SLE items screened in Colodro-Conde *et al*. compared to our study. Although questions about dependent SLE (DSLE, mean = 0.4 SLE) represented over 28% of the TSLE-items reported in our study, the main effect of DSLE explained approximately 93% of the amount of variance explained by TSLE (*full cohort:* variance explained = 4.56%, β = 0.212, s.e. = 0.014, *p* = 1.73 × 10^−54^; *men:* 3.74%, β = 0.193, s.e. = 0.021, *p* = 9.66 × 10^−21^; *women:* 5.07%, β = 0.225, s.e. = 0.018, *p* = 8.09 × 10^−35^; Figure 1b). Independent SLE (ISLE, mean = 0.85 SLE), which represented over 69% of TSLE-items, explained approximately 57% of the amount of variance explained by TSLE (*full cohort:* variance explained = 2.80%, β = 0.167, s.e. = 0.014, *p* = 1.32 × 10^−33^; *men*: 2.44%, β = 0.156, s.e. = 0.022, *p* = 2.88 × 10^−13^; *women:* 3.02%, β = 0.174, s.e. = 0.018, *p* = 5.20 × 10^−22^; Figure 1b). To explore the contribution from each measure, we combined DSLE and ISLE together in a single model. DSLE explained 3.34% of the variance of depressive score compared to 1.45% of the variance being explained by ISLE, suggesting that DSLE have a greater effect on liability to depressive symptoms than ISLE.

**Figure 1.**
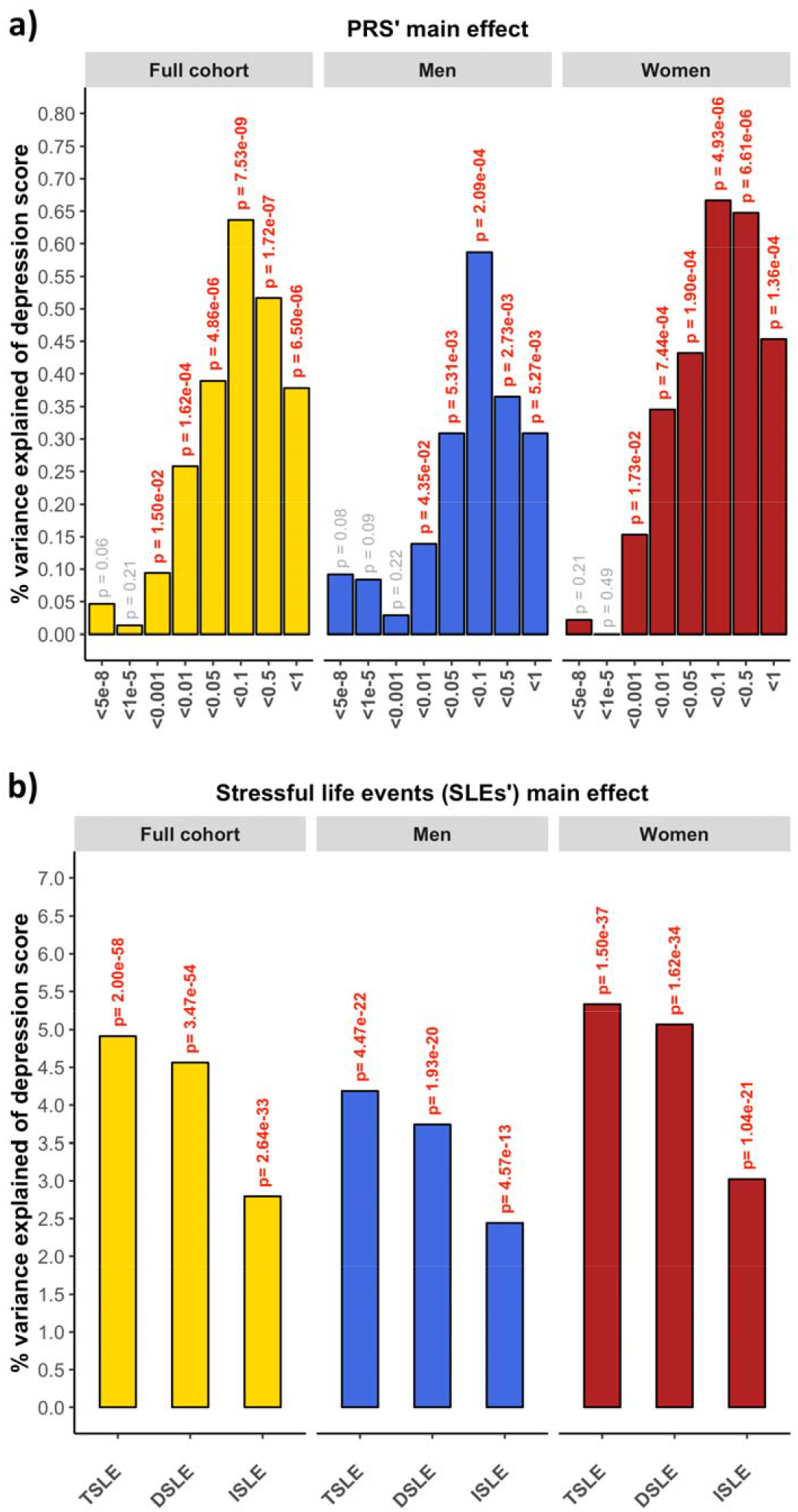
**a)** Association between polygenic risk scores (PRS) and depression score (main effects, one-sided tests). PRS were generated at 8 *p*-threshold levels using summary statistics from the Psychiatric Genetic Consortium MDD GWAS (released July 2016) with the exclusion of Generation Scotland participants. The depression score was derived from The General Health Questionnaire. The Y-axis represents the % of variance of depression score explained by PRS main effects. The full cohort (yellow) was split into men (blue) and women (red). In Colodro-Conde *et al*., PRS for MDD significantly explained up to 0.46% of depression score in their sample (∼0.39% in women and ∼0.70% in men), **b)** Association between reported number of SLE and depression score (main effect, one-sided tests, results expressed in % of depression score explained). SLE were self-reported through a brief life-events questionnaire based on the List of Threating Experiences and categorized into: total number of SLE reported (TSLE), “dependent” SLE (DSLE) or “independent” SLE (ISLE). The full cohort (yellow) was split into men (blue) and women (red). In Colodro-Conde *et al*., “personal” SLE significantly explained up to 12.9% of depression score variance in their sample (∼11.5% in women and ∼16% in men).

A *diathesis-stress* model for depression was tested to assess GxE effects. We detected significant, albeit weak, GxE effects on depression score (Figure 2). The PRS interaction with TSLE was nominally significant in the full cohort (β = 0.028, s.e. = 0.014, R^2^ = 0.08%, *p* = 0.049) and slightly stronger in women (β = 0.044, s.e. = 0.018, R^2^ = 0.19%, *p* = 0.017; Figure 2a), compared to men in which the effect was not significant (β = 0.039, s.e. = 0.022, R^2^ = 0.15%, *p* = 0.07). However, these results did not survive correction for multiple testing (*p* > 2.08 × 10^−3^).

**Figure 2.**
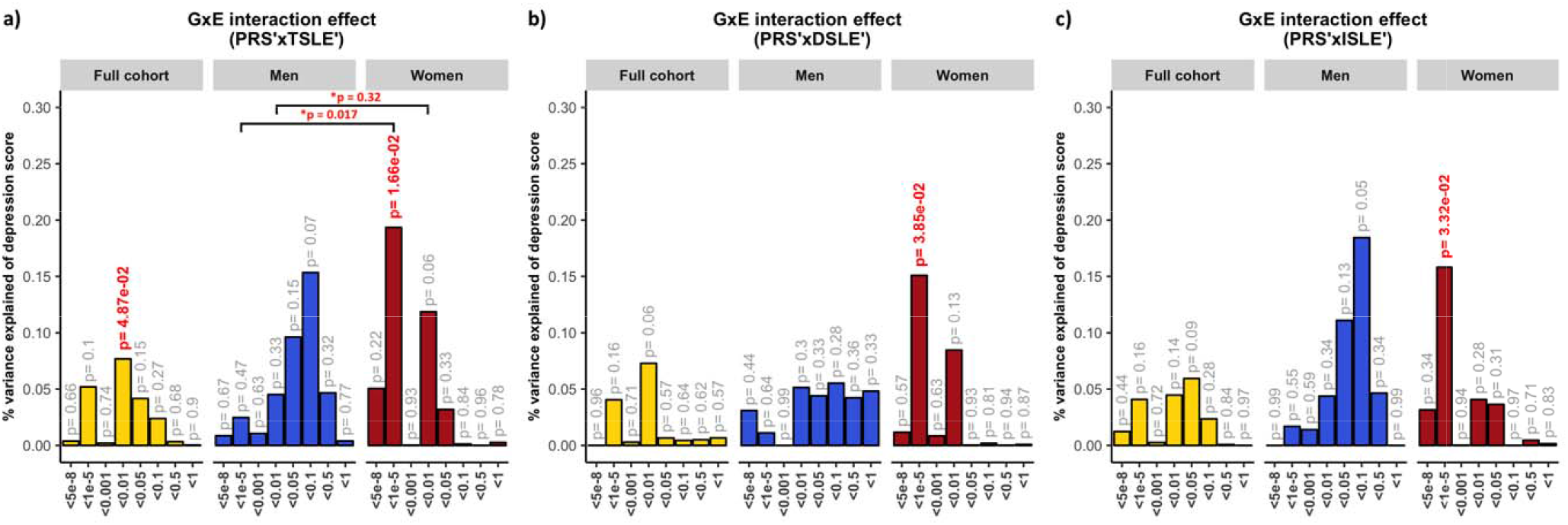
Association between GxE effect and depression score. The results represent percentage of depression score explained by the interaction term (two-sided tests) fitted in linear mixed models to empirically test the *diathesis-stress* model. Red numbers show significant interactions *p*-values. *Shows significance of difference between sexes when comparing the size of the estimated GxE effects. The full cohort (yellow) was split into men (blue) and women (red). PRS were generated at 8 *p*-threshold levels using summary statistics from the Psychiatric Genetic Consortium MDD GWAS (released July 2016) with the exclusion of Generation Scotland participants. The interaction effect was tested with **a)** the number of SLE reported (TSLE), **b)** “dependent” SLE (DSLE) and **c)** “independent” SLE (ISLE). In Colodro-Conde *et al*., the variance of depression score explained in their sample by GxE was 0.12% (*p* = 7 × 10^−3^). GxE were also significant in women (*p* = 2 × 10^−3^) explaining up to 0.25% of depression score variation, but not in men (*p* = 0.059; R^2^ = 0.17%; negative/protective effect on depression score).

The best-fit threshold was much lower in women (*p*-threshold = 1 × 10^−5^) compared to the full sample (*p*-threshold = 0.01). The size of GxE across sexes at *p*-threshold = 1 × 10^−5^ were significantly different (GxE*sex *p* = 0.017), but not at the best *p*-threshold in the full cohort (*p*-threshold = 0.01, GxE*sex *p* = 0.32; Figure 2a). In women, GxE effect with DSLE predicted depression score (*p*-threshold = 1 × 10^−5^; β = 0.039, s.e. = 0.019, R^2^ = 0.15%, *p* = 0.038; Figure 2b and Supplementary Figure 2a), as did the GxE effect with ISLE (*p*-threshold = 1 × 10^−5^; β = 0.040, s.e. = 0.019, R^2^ = 0.16%, *p* = 0.033; Figure 2c and Supplementary Figure 2b). No significant interaction was detected in men (best-fit *p*-threshold = 0.1) with either TSLE (β = 0.039, s.e. = 0.022, R^2^ = 0.15%, *p* = 0.072; Figure 2a), DSLE (β = 0.024, s.e. = 0.022, R^2^ = 0.06%, *p* = 0.28; Figure 2b) or ISLE (β = 0.043, s.e. = 0.022, R^2^ = 0.18%, *p* = 0.055; Figure 2c).

To examine these results further and investigate the *diathesis* effect at different levels of stress, nominally significant GxE were plotted between PRS and categories of SLE (i.e, “none”, “low” and “high” SLE reported; Figure 3). Examining the interaction found at the full cohort (PRS at PGC-MDD GWAS *p*-threshold = 0.01), we detected a significant direct *diathesis* effect on the risk of depressive symptoms in those participants reporting SLE, with a higher risk when greater numbers of SLE were reported (“low” number of SLE reported: PRS’ β = 0.043, s.e. = 0.021, *p* = 0.039; “high” number of SLE reported: PRS’ β = 0.142, s.e. = 0.039, *p* = 2.86 × 10^−4^; see Table 1 and Figure 3a). Whereas, in participants who reported no SLE over the preceding 6 months, the risk of depressive symptoms was the same regardless of their *diathesis* risk (“none” SLE reported: PRS’ β = 0.021, s.e. = 0.022, *p* = 0.339). Stratifying these results by sex, we found the same pattern as in the full cohort in women (“none”: *p* = 0.687; “low”: *p* = 0.023; “high”: *p* = 2 × 10^−3^), but not in men (“none”: *p* = 0.307; “low”: *p* = 728; “high”: *p* = 0.053; see Table 1 and Figure 3a). However, the lack of significant *diathesis* effect in men may be due to their lower sample size and its corresponding reduced power.

**Table 1.**
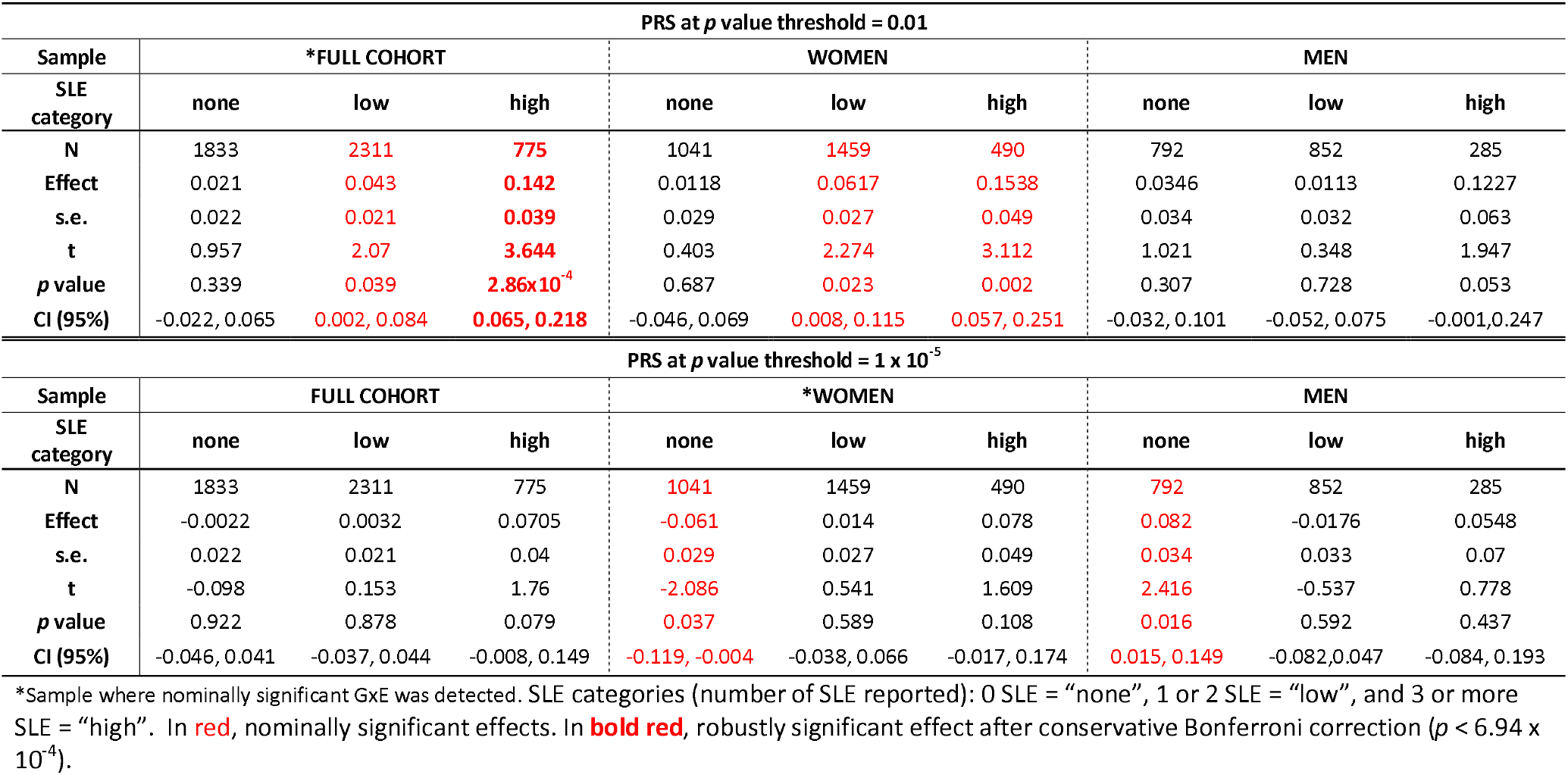
Diathesis effect on depression score under SLE categories. Reported values at p-thresholds where nominally significant GxE effects were detected.

**Figure 3.**
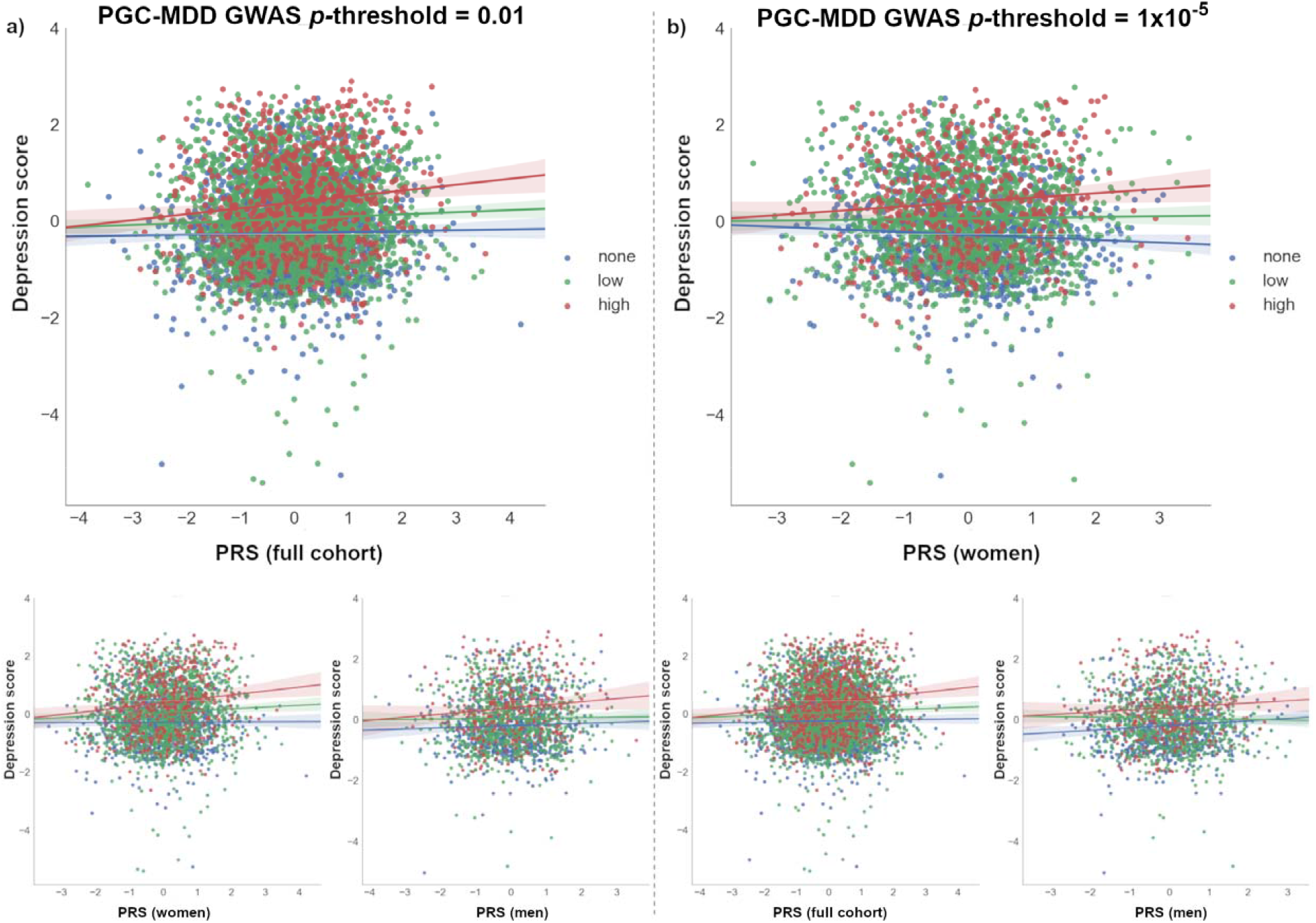
Scatterplot of *diathesis-stress* interactions on depression score. Interactions with PRS at which nominally significant GxE effects were detected in **a)** full cohort (*p*-threshold = 0.01) and **b)** in women (*p*-threshold = 1 × 10^−5^) are shown. At bottom, the remaining samples (i.e., *full cohort, women* or *men*) at same *p*-threshold are shown for comparison. The X-axis represents the direct effect of PRS (standard deviation from the mean) based on **a)** *p*-threshold = 0.01 and **b)** *p*-threshold = 1 × 10^−5^ using the total number of SLE reported by each participant (dot) as environmental exposures at three SLE levels represented by colours. Blue: 0 SLE, “no stress”, n = 1 833/1 041/792; green: 1 or 2 SLE, “low stress”, n = 2 311/1 459/852; red: 3 or more SLE, “high stress”, n = 775/490/285; in the full cohort, women and men, respectively. Y-axis reflects the depression score standardized to mean of 0 and standard deviation of 1. Lines represent the increment in risk of depression under a certain degree of “stress” dependent on a genetic predisposition (= *diathesis*).

Examining the interaction with PRS (at PGC-MDD GWAS *p*-threshold = 1 × 10^−5^) with which a significant interaction was detected in women, we only detected a significant *diathesis* effect on depression score when stratifying by sex in those participants who did not reported SLE over the last 6 months (see Table 1). The *diathesis* effect was positive in men (PRS’ β = 0.082, s.e. = 0.034, *p* = 0.016, R^2^ = 0.7%; Figure 3b), consistent with the contribution of risk alleles. Conversely, the *diathesis* effect was negative in women (PRS’ β = −0.061, s.e. = 0.029, *p* = 0.037, R^2^ = 0.4%; Figure 3b), suggesting a protective effect of increasing PRS in those women reporting no SLE, and consistent with the contribution of alleles to individual sensitivity to both positive and negative environmental effects (i.e. “plasticity alleles” rather than “risk alleles”)^53,54^. This PRS accounted for the effect of just 34 SNPs, and the size of its GxE across sexes were significantly different (GxE*sex *p* = 0.017; Figure 2a), supporting possible differences in the underlying stress-response mechanisms between women and men.

## DISCUSSION

The findings reported in this study support those from Colodro-Conde *et al*.^23^, in an independent sample of similar sample size and study design, and also supports possible sex-specific differences in the effect of genetic risk of MDD in response to SLE. Both Colodro-Conde *et al*. and our study suggest that individuals with an inherent genetic predisposition to MDD, reporting high number of recent SLE, are at additional risk of depressive symptoms due to GxE effects, thus validating the *diathesis-stress* theory. We identified nominally significant GxE effects in liability to depression at the population level (*p* = 0.049) and in women (*p* = 0.017), but not in men (*p* = 0.072). However, these interactions did not survive multiple testing correction (*p* > 2.08 × 10^−3^) and the power of both studies to draw robust conclusions remains limited^55^. With increased power these studies could determine more accurately both the presence and magnitude of a GxE effect in depression. To better understand the effect of PRS at different levels of exposure to stress, we examined the nominally significant interactions detected in the full sample by categorizing participants on three groups based on the number of SLE reported (i.e. “none”, “low” or “high”). We detected a significant *diathesis* effect on risk of depression only in those participants reporting SLE, but not in those participants that reported no SLE over the last 6 month. Furthermore, the *diathesis* effect was stronger on those participants reporting a “high” number of SLE (β = 0.142, *p* = 2.86 × 10^−4^) compared to those participants reporting a “low” number of SLE (β = 0.043, *p* = 0.039). The former effect was robustly significant and survived a conservative Bonferroni correction to adjust for multiple testing (*p* < 6.94 × 10^−4^). This finding corroborates the *diathesis-stress* model for depression and supports Colodro-Conde *et al*. results using an independent sample.

To investigate the relative contribution of the GxE to the variance of depression, we examined in the full cohort the total variance of depression score explained by the PRS main effect and the significant GxE effect jointly. Together, they explained 0.34% of the variance, of which 0.07% of the variance of the depression score was attributed to the GxE effect (*p*-threshold = 0.01; PRS *p* = 1.19 × 10^−4^, GxE *p* = 0.049; both derived from the full diathesis-model with TSLE). This is lower than the proportion of variance attributed to common SNPs (8.9%) in the full PGC-MDD analysis^16^. As Colodro-Conde *et al*. noted, this result aligns with estimates from experimental organisms suggesting that around 20% of the heritability may be typically attributed to the effects of GxE^56^, although it is inconsistent with the majority of human traits with the potential exception of depression^57^.

Consistent with PRS predicting “personal” SLE in Colodro-Conde *et al*., PRS for MDD predicted SLE in our study (see Supplementary Figure 1), although not at the *p*-threshold at which significant GxE effects were detected. Genetic factors predisposing to MDD may contribute to individuals exposing themselves to, or showing an increased reporting of, SLE via behavioural or personality traits^58,59^. Such genetic mediation of the association between depression and SLE would disclose a gene-environment correlation (i.e. genetic effects on the probability of undergoing a SLE) that hinders to interpret our findings as pure GxE effects^60,61^. To address this limitation and assess this aspect, following Colodro-Conde *et al*., we split the 12-items TSLE measure into SLE that are either potentially “dependent” on a participant’s own behaviour (DSLE; therefore, potentially driven by genetic factors) or not (“independent” SLE; ISLE)^46,49^. DSLE are reported to be more heritable and have stronger associations with MDD than ISLE^49,62,63^. In our sample, reporting DSLE is significantly heritable (h^2^_SNP_ = 0.131, s.e. = 0.071, *p* = 0.029), supporting a genetic mediation of the association, whereas reporting ISLE is not significantly heritable (h^2^_SNP_ = 0.000, s.e. = 0.072, *p* = 0.5). Nominally significant GxE effects were seen in women for both DSLE and ISLE, suggesting that both GxE and gene-environment correlation co-occur. Colodro-Conde *et al*. did not identify significant GxE using independent SLE as the exposure.

Between-sex differences on stress response could help to explain previous differences seen between sexes in depression such as those in associated risk (i.e. approximately 1.5 - 2-fold higher in women), symptoms reported and/or coping strategies (e.g., whereas women tend to cope through verbal and emotional strategies, men tend to cope by doing sport and consuming alcohol)^64,68^. This also aligns with an increased risk associated with a lack of social support seen in women compared to men^23^. Furthermore, although we do not know whether participants experienced recent events with positive effects, we saw a protective effect in those women who did not experienced recent SLE (*p* = 0.037), suggesting that some genetic variants associated with MDD may operate as “plasticity alleles” and not just as “risk alleles^-53,54^. This effect was neutralized in the full cohort due to an opposite effect in men (*p* = 0.016), but it is supported by previous protective effects reported when using a serotoninergic multilocus profile score and absence of SLE in young women^69^. These findings would be consistent with a differential-susceptibility model^70,71^ of depression, also suggested by the interaction effects seen between the serotonin transporter linked promoter region gene (5-HTTLPR) locus and family support and liability to adolescent depression in boys^72^. However, our results and the examples given are only nominally significant and will require replication in larger samples. Robustly identified sex-specific differences in genetic stress-response could improve personalized treatments and therapies such as better coping strategies.

There are notable differences between our study and Colodro-Conde *et al*. to consider before accepting our findings as a replication of Colodro-Conde *et al*. results. First, differences in PRS profiling may have affected replication power. We used the same equivalent PGC-MDD2 GWAS as discovery sample. However, whereas Colodro-Conde *et al*. generated PRS in their target sample containing over 9.5M imputed SNP, in this study we generated PRS in a target sample of over 560K genotyped SNPs (see Supplementary table 1 for comparison). This potentially results in a less informative PRS in our study, with less predictive power, although the variance explained by our PRS was slightly larger (0.64% vs. 0.46%). The size of the discovery sample is key to constructing an accurate predictive PRS, but to exploit the most of the variants available may be an asset^55^. Secondly, different screening tools were used to measure both current depression and recent environmental stressors across the two studies. Both studies transformed their data, using item response theory or by log-transformation, to improve the data distribution. However, neither study used depression scores that were normally distributed. The scale of the instruments used and their corresponding parameterization to test an interaction could have a direct effect on the size and significance of their interaction^56,73^; so findings from GxE must be taken with caution. Furthermore, although both screening methods have been validated and applied to detect depressive symptoms, different questions may cover and emphasise different features of the illness, which may result in different outputs. The same applies to the measurement of environmental stressors in the two studies. Both covering of a longer time-period and upweighting by “dependent” SLE items may explain the increased explanatory power of “personal” SLE (12.9%) in Colodro-Conde *et al*. to predict depression score compared to our “total” SLE measure (4.91%). Finally, the unmeasured aspects of the exposure to SLE or its impact may also contribute to lack of stronger replication and positive findings.

In conclusion, despite differences in the measures used across studies, we saw concordance and similar patterns between our results and those of Colodro-Conde *et al*.^23^. Our findings are consistent with Colodro-Conde *et al*. and, therefore, add validity to the *diathesis-stress* theory for depression. Empirically demonstrating the *diathesis-stress* theory for depression would validate recent^20-22,24^ and future studies using a genome-wide approach to identify genetic mechanisms and interactive pathways involved in GxE underpinning the causative effect of “stress” in the development of depressive symptoms and mental illness in general. This study adds to our understanding of gene-by-environment interactions, although larger samples will be required to confirm differences in *diathesis-stress* effects between women and men.

## Supporting information

## ACKNOWLEDGMENTS

The 1^st^ author AAS is funded by the University of Edinburgh (www.ed.ac.uk) and MRC for his PhD study at the University of Edinburgh Institute of Genetics and Molecular Medicine (www.ed.ac.uk/igmm). DJM acknowledges the financial support of NHS Research Scotland (NRS) through NHS Lothian. MA is supported by STRADL through a Wellcome Trust Strategic Award (reference 104036/Z/14/Z). Generation Scotland received core support from the Chief Scientist Office of the Scottish Government Health Directorates [CZD/16/6] and the Scottish Funding Council [HR03006]. The genotyping of the GS:SFHS samples was carried out by the Genetics Core Laboratory at the Wellcome Trust Clinical Research Facility, Edinburgh, Scotland and was funded by the Medical Research Council UK and the Wellcome Trust (Wellcome Trust Strategic Award “STratifying Resilience and Depression Longitudinally” (STRADL) Reference 104036/Z/14/Z). The Major Depressive Disorder Working Group of the Psychiatric Genomics Consortium depends on the contributions of many parties.

## FINANCIAL DISCLOSURE

The authors declare no conflict of interest.

